# Mutational impact of chronic alcohol use on stem cells in cirrhotic liver

**DOI:** 10.1101/698894

**Authors:** Myrthe Jager, Ewart Kuijk, Ruby Lieshout, Mauro D. Locati, Nicolle Besselink, Bastiaan van der Roest, Roel Janssen, Sander Boymans, Jeroen de Jonge, Jan N.M. IJzermans, Michael Doukas, Monique M.A. Verstegen, Ruben van Boxtel, Luc J.W. van der Laan, Edwin Cuppen

**Affiliations:** Center for Molecular Medicine and Oncode Institute, University Medical Center Utrecht, Utrecht University, Heidelberglaan 100, 3584 CX Utrecht, The Netherlands; Department of Surgery, Erasmus MC - University Medical Center, Wytemaweg 80, 3015 CN Rotterdam, The Netherlands; Department of Pathology, Erasmus MC - University Medical Center, Wytemaweg 80, 3015 CN Rotterdam, The Netherlands

**Author notes:** Oncode Institute and Princess Máxima Center for Pediatric Oncology, 3584 CT Utrecht, The Netherlands.

## Abstract

Excessive alcohol consumption increases the risk of developing liver cancer, but the mechanism through which alcohol drives carcinogenesis is as yet unknown. Here, we determined the mutational consequences of chronic alcohol use on the genome of human liver stem cells prior to cancer development. No change in base substitution rate or spectrum could be detected. Analysis of the trunk mutations in an alcohol-related liver tumor by multi-site whole-genome sequencing confirms the absence of specific alcohol-induced mutational signatures driving the development of liver cancer. However, we did identify an enrichment of nonsynonymous base substitutions in cancer genes in stem cells of the cirrhotic livers, such as recurrent nonsense mutations in *PTPRK* that disturb Epidermal Growth Factor (EGF)-signaling. Our results thus suggest that chronic alcohol use does not contribute to carcinogenesis through altered mutagenicity, but instead induces microenvironment changes which provide a ‘fertile ground’ for selection of cells with oncogenic mutations.

## INTRODUCTION

Alcohol consumption is an important risk factor for the development of various cancer types, including hepatocellular carcinoma (HCC), and causes an estimated 400,000 cancer-related deaths worldwide each year^1–3^. In spite of the clear link between alcohol intake and tumorigenesis, the underlying mechanism remains debated and mainly revolves around two hypotheses. The first hypothesis suggests that alcohol consumption may contribute to the development of cancer through an increased mutation accumulation in the genome^4^. Consistently, the first metabolite of ethanol, acetaldehyde, is highly carcinogenic^5–7^ and can also contribute to the formation of mutagenic reactive oxygen species (ROS)^8–11^. Analysis of a large number of tumor exomes and genomes showed that alcohol intake is associated with an increased mutation load and different mutational characteristics^12–15^. The second hypothesis suggests that an alcohol-induced change of microenvironment is an essential driver for tumorigenesis by providing a fertile ground for cells with oncogenic mutations^16–18^. Indeed, development of HCC is preceded by chronic inflammation and cirrhosis in about 80% of patients and this cell-extrinsic damage appears a prerequisite for the formation of the majority of liver cancers^18–20^. Additionally, Hepatitis C Virus (HCV)-induced cirrhotic livers show an increase in the number and size of clonal patches with mutations in genes that are frequently mutated in HCC^21^. Alcohol use itself has been associated with an increased number of cancer-stem-cell-like epithelial cell adhesion molecule (EpCAM)-positive cells in the cirrhotic liver^22^, which may be driven by epithelial to mesenchymal transition through activation of the Wnt pathway^23^, confirming that cellular composition changes can be induced by alcohol use. Yet, it is still uncertain whether an altered cellular environment is sufficient to drive the development of cancer, or whether an increase in the mutation load is also required. The here mentioned hypotheses are thus not mutually exclusive.

We have demonstrated previously that mutations accumulate linearly with age in liver adult stem cells (ASCs) of healthy individuals, without controlling for lifestyle^24,25^. Stem cells are believed to be an important cell-of-origin for several cancer types, including liver cancer^18,26–28^, although liver cancer can also originate in differentiated cells^29–31^. Here, we studied the accumulation of mutations in ASCs from non-cancerous, cirrhotic livers of patients with a history of chronic alcohol use and compared these to the mutational patterns of healthy liver donors and to mutations that accumulated in the most recent common ancestor (MRCA) cell of an alcohol-related HCC.

## RESULTS

### Mutation load similar in alcoholic liver

We sequenced the genomes of eight independent clonal organoid cultures derived from biopsies of five non-cancerous, cirrhotic livers from patients with a known history of chronic alcohol intake who were undergoing a liver transplantation (further referred to as ‘alcoholic livers’; Supplementary Table 1). To gain insight into the mutational consequences of chronic alcohol consumption, the somatic mutation catalogs of alcoholic livers were compared to those obtained previously from whole genome sequencing (WGS) data of five healthy liver donors (further referred to as ‘healthy livers’)^24^. To increase the number of healthy liver donors and to obtain age-matched healthy controls, five clonal liver organoid cultures derived from four additional healthy liver donors with ages ranging from 24 to 68 years were included in the analyses (Supplementary Table 1).

In total, we identified 42,093 base substitutions, 1,931 indels, and 5 copy number alterations (CNAs) (Fig. 1; Supplementary Fig. 1; Supplementary Table 1). Consistent with previous observations^24^, there is a positive relationship between somatic base substitutions and age in healthy liver ASCs (two-tailed *t*-test, linear mixed model, *P* < 0.05; Fig. 1a). Healthy liver ASCs acquired ∼39.4 (95% confidence interval (95% CI): 30.5 - 48.3) somatic base substitutions each year. The mutation load in alcoholic liver ASCs (Fig. 1a) was similar to, and within the 95% CI of the slope estimate of, age-matched healthy liver ASCs. Alcohol consumption did not affect the number of tandem base substitutions acquired in the genomes of liver ASCs either (Supplementary Fig. 1a). Furthermore, indels also accumulated with age at a comparable rate in healthy and alcoholic liver ASCs (two-tailed *t*-test, linear mixed model, *P* < 0.05; Supplementary Fig. 1b). Finally, a minority of the healthy liver ASCs and none of the assessed alcoholic liver ASCs acquired a CNA, although few CNAs were detected (Supplementary Table 1)^24^. At the chromosomal level we observed trisomy 22 in a healthy liver ASC from a 68-year-old healthy female donor and chromosome Y gain in an alcoholic liver ASC from a 67-year-old male donor (Supplementary Fig. 1c). This suggests that aneuploidies may occur in liver ASCs of older individuals, but this seems to be unrelated to alcohol consumption. Taken together, these results strongly suggest that the induction of HCC by chronic alcohol consumption is not caused by an altered base substitution, indel, or CNA accumulation in liver ASCs prior to oncogenesis.

**Fig. 1.**
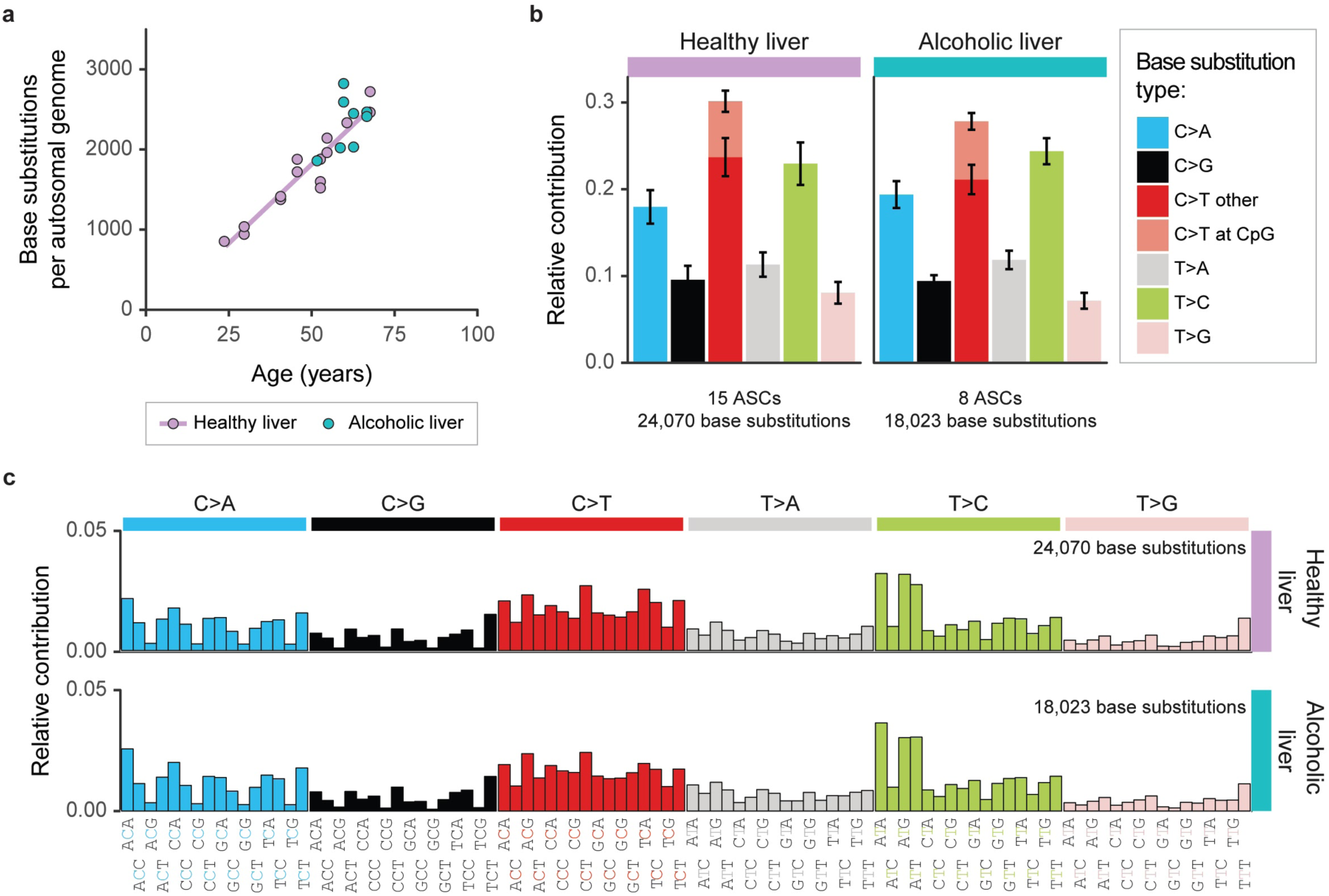
Somatic base substitutions in healthy and alcoholic liver stem cells. **a** Number of somatic base substitutions in the autosomal genomes of 15 healthy and 8 alcoholic liver stem cells of 9 and 5 donors, respectively. Each stem cell is represented by a data point. A linear accumulation of base substitutions with age was observed in healthy liver, indicated by the purple trendline. **b** Mean relative contribution of the base substitution types to the mutation spectra of healthy and alcoholic liver ASCs. Error bars represent standard deviation. **c** Mean relative contribution of 96 context-dependent base substitution types to the mutational profiles of healthy and alcoholic liver ASCs.

### Mutation type similar in alcoholic liver

It has been shown that genome-wide patterns of base substitutions reflect past activity of mutational processes in cells^32^. Previously, alcohol consumption was reported to be associated with a modest increase of Catalogue Of Somatic Mutations In Cancer (COSMIC) signature 16 mutations, which is characterized by T:A>C:G mutations^32^, in esophageal and liver cancer^12–15^. To identify if excessive alcohol consumption changed the mutational profiles in non-cancerous liver ASCs, we performed in-depth mutational analyses. The mutational profiles of healthy liver ASCs were characterized by a high contribution of C:G>A:T, C:G>T:A, and T:A>C:G mutations (Fig. 1b-c; Supplementary Fig. 2). The mutational profiles of alcoholic liver ASCs were highly similar to the mutational profiles of healthy liver ASCs (cosine similarity = 0.99), indicating that chronic alcohol use does not alter the mutational processes in liver ASCs.

**Fig. 2.**
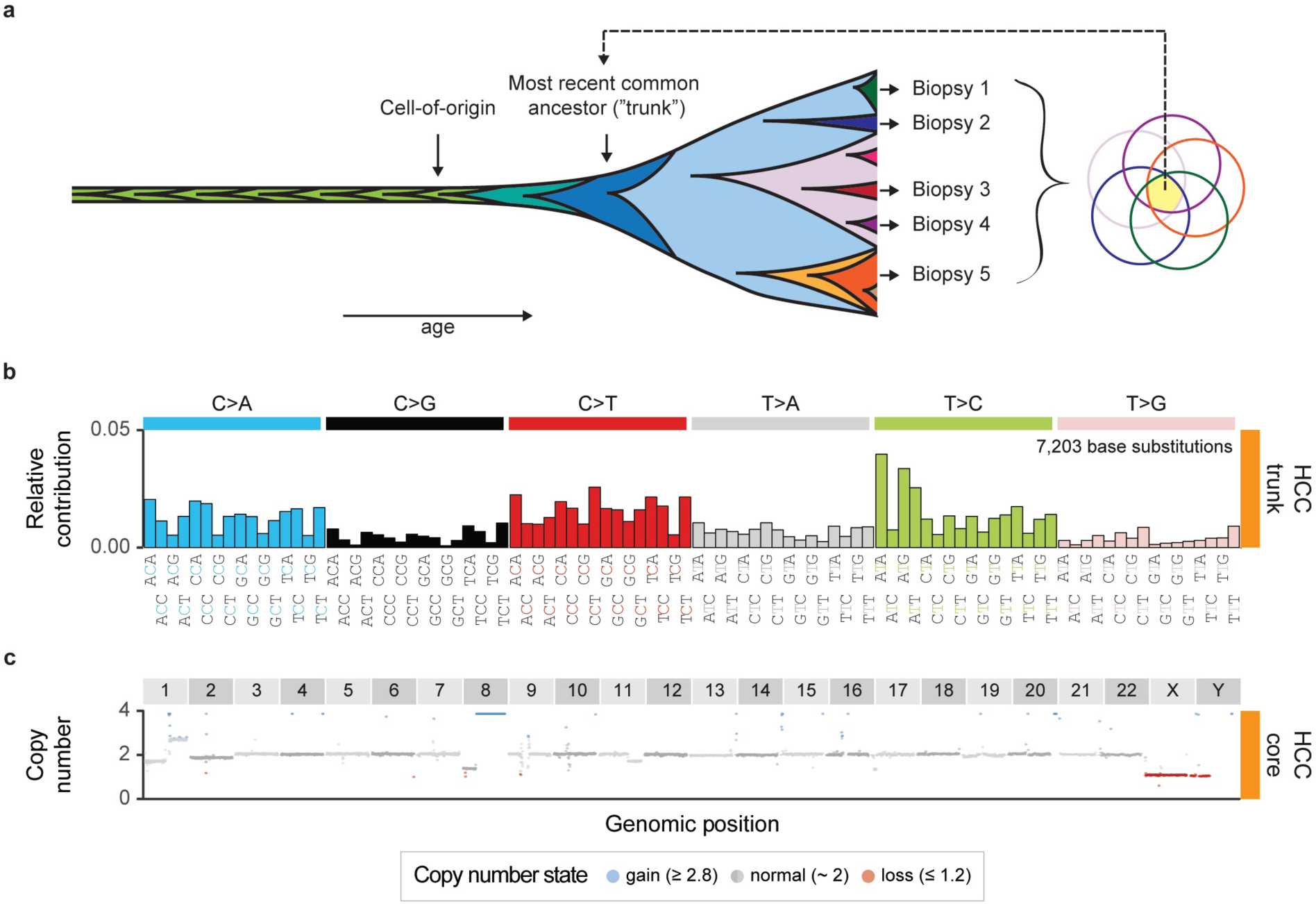
Somatic mutations in the most recent common ancestor of an HCC in a patient with a history of chronic alcohol intake. **a** Mutations that accumulated in the most recent common ancestor (“trunk”) can be identified, by determining the mutations that are shared by multiple biopsies (in yellow). Five biopsies across a 13cm HCC were sequenced to identify the trunk mutations. **b** Relative contribution of 96 context-dependent base substitution types to the mutational profile of the recent common ancestor of an HCC. **c** Genomic copy number profile of one of the HCC biopsies (HCC-core), which is representative for all biopsies (Supplementary Fig. 4c).

To determine whether chronic alcohol use changed the contribution of the known COSMIC mutational signatures^32–34^, we calculated the contribution of these signatures to the mutational profiles of all ASCs and, subsequently, performed a bootstrap resampling method to identify potential significant differences between healthy and alcoholic liver ASCs, similar as described in Zou *et al*. ^35^. COSMIC signatures 5 and 40 could explain the majority of the accumulated base substitutions in both healthy and alcoholic liver (Supplementary Fig. 3). However, we did not observe a significant change in signature contributions between alcoholic liver ASCs and healthy liver ASCs (bootstrap resampling method, see Methods; Supplementary Fig. 3).

**Fig. 3.**
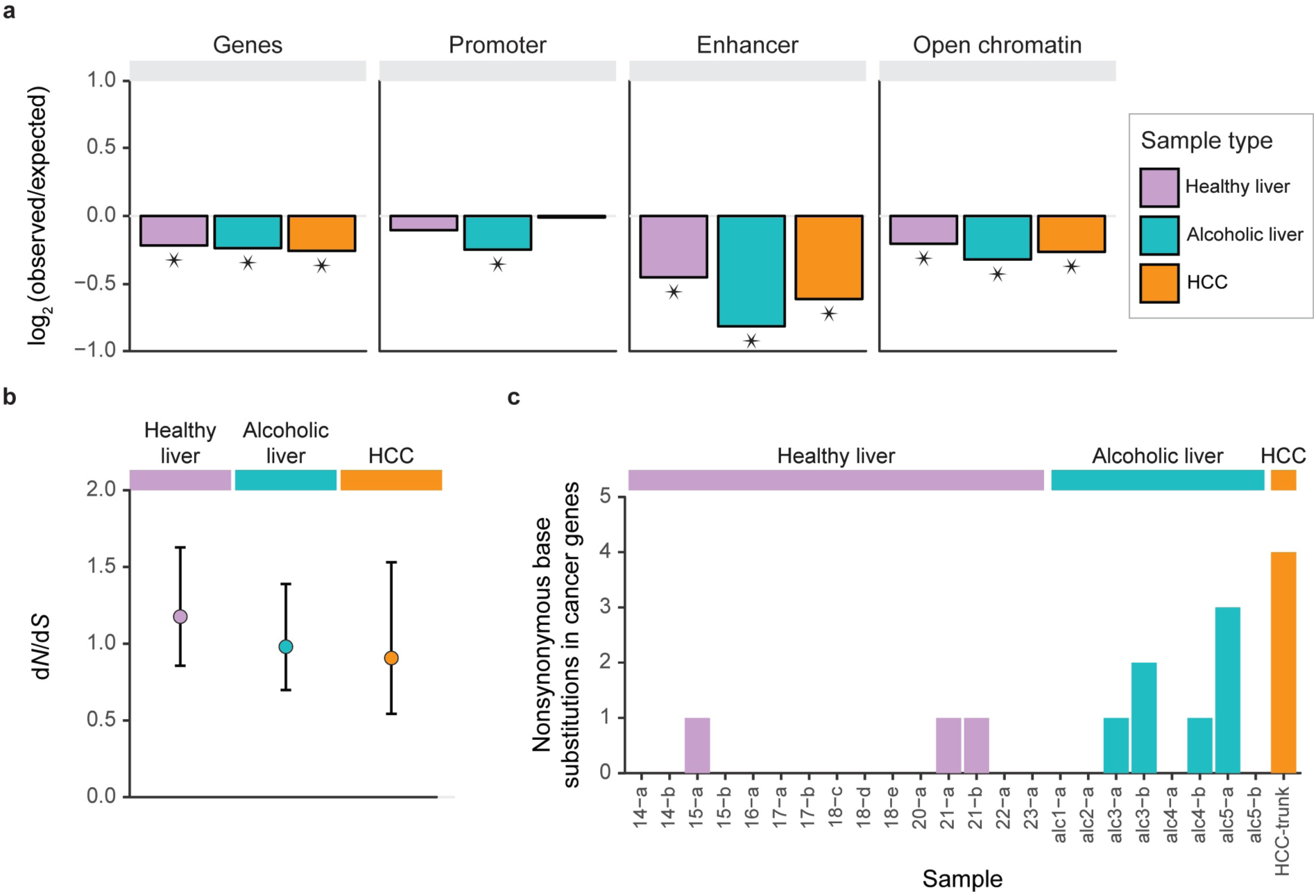
Genomic distribution of the somatic base substitutions acquired in healthy liver stem cells, alcoholic liver stem cells, and the most recent common ancestor of an HCC. **a** The effect size of the depletion of somatic base substitutions in genes, promoters, enhancers, and open chromatin regions. Asterisks indicate significant depletion. **b** d*N*/d*S* of the somatic base substitutions in genes in the indicated sample types. Data points represent the Maximum-likelihood estimates and error bars represent the 95% confidence intervals. **c** Number of nonsynonymous base substitutions in cancer genes in each indicated sample.

A possible explanation for the absence of a correlation between alcohol consumption and mutational patterns is that the cells that we have sequenced are too early in the precancerous state. Therefore, we also sequenced five biopsies across a 13 cm HCC of a 60-year-old male donor with a history of chronic alcohol use and identified mutations that were shared by all biopsies (Fig. 2a; Supplementary Fig. 4). This approach allowed for the identification of all mutations in the MRCA and thus provided insight into the mutational process that had been active prior to tumor formation and in the early to intermediate stages of tumor development (Fig. 2a). As a control sample, we sequenced a non-tumorous biopsy adjacent to the tumor, to identify and exclude germline mutations. In total, we identified 19,200 unique somatic base substitutions across all five HCC biopsies (Supplementary Table 2; Supplementary Fig. 4b).

**Fig. 4.**
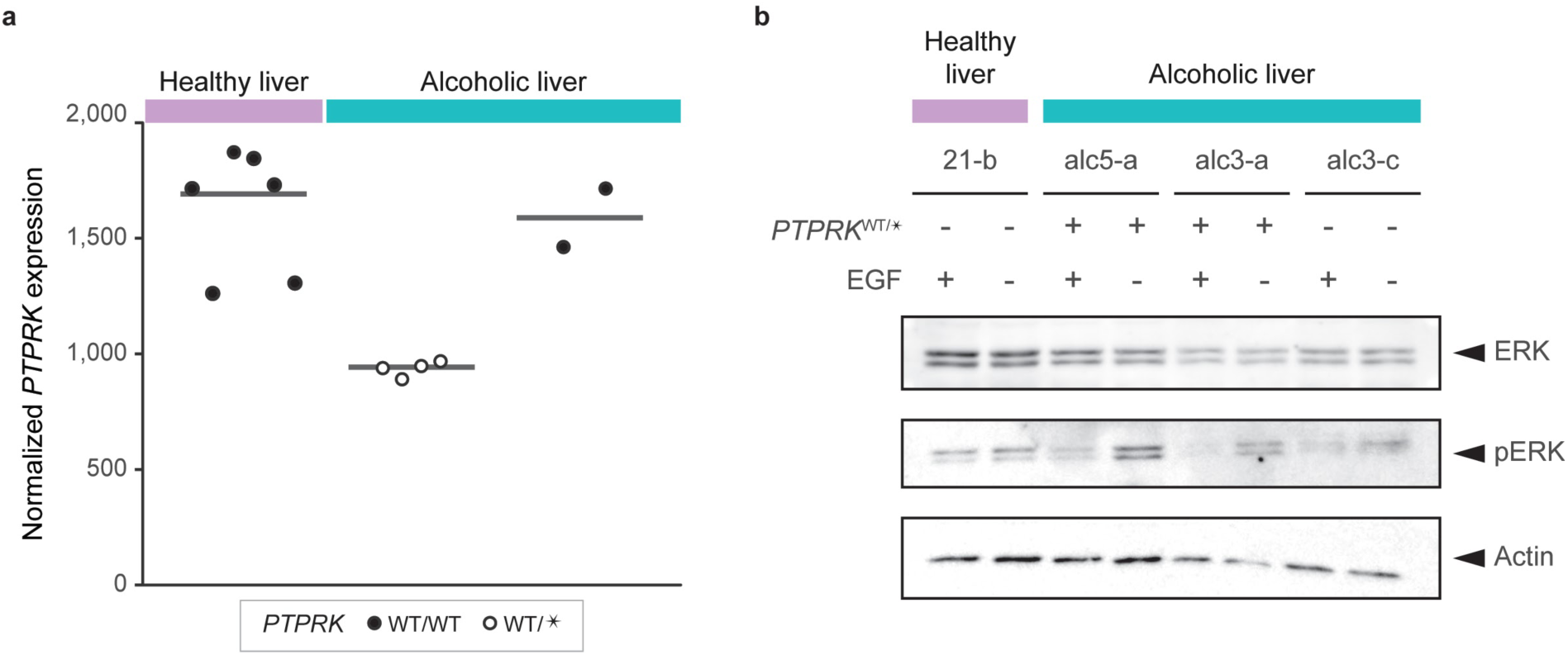
Functional consequences of nonsense *PTPRK* base substitutions. **a** Normalized *PTPRK* mRNA expression in alcoholic *PTPRK*^WT/*^, healthy *PTPRK*^WT/WT^ and alcoholic *PTPRK*^WT/WT^ liver organoid cultures. Normalized counts were calculated for duplicate measures of 2 alcoholic *PTPRK*^WT/*^, 3 healthy *PTPRK*^WT/WT^, and 1 alcoholic *PTPRK*^WT/WT^ organoid cultures. Each data point represents a single measurement. Lines indicate median *PTPRK* expression per sample type. WT = wildtype, * = nonsense base substitution. **b** Western blot of ERK, pERK and actin in organoid cultures of *PTPRK*^WT/*^ livers and healthy and alcoholic *PTPRK*^WT/WT^ liver cultures with and without EGF in the culturing medium.

Analysis of the base substitutions shared by all biopsies (trunk mutations) revealed that the MRCA of these biopsies accumulated 7,203 base substitutions (Supplementary Table 2; Supplementary Fig. 4b). The mutational profile of the trunk mutations in the HCC was highly similar to healthy and alcoholic liver ASCs (Fig. 2b; cosine similarity = 0.97 and 0.98, respectively). These results were in line with our initial observations in ASCs that alcohol itself does not introduce specific mutations in the genome of liver cells. The high mutation load suggests that the MRCA already evolved significantly from the cell-of-origin (Fig. 2a) and that a clonal sweep occurred after a substantial amount of mutations already accumulated. Consistently, the MRCA already acquired two CNAs (Supplementary Table 3) and several chromosomal aneuploidies (Fig. 2c; Supplementary Fig. 4c).

### Cancer driver mutations

Previously, alcohol intake has been shown to accelerate the expansion of clones with cancer driver mutations in the esophagus^36^. To identify whether chronic alcohol consumption induces similar changes in the selection of liver cells, we analyzed the genomic distribution of the acquired base substitutions. If chronic alcohol use would affect cellular selection, the frequency of somatic mutations in active functional genomic elements would differ between alcoholic and healthy liver. Base substitutions were, however, depleted to a similar extend in regions such as genes and enhancers in healthy liver ASCs and alcoholic liver ASCs (Fig. 3a). Furthermore, unlike previous observations^37^, we did not observe an enrichment of base substitutions in H3K36me3 regions, associated with active transcription^38,39^, in alcoholic liver ASCs in comparison to healthy liver ASCs (Supplementary Fig. 5). The normalized ratio of nonsynonymous to synonymous base substitutions (d*N*/d*S*) was also ∼1 in all assessed cell types (Fig. 3b). Taken together, these results suggest that there is no general change in selection against more deleterious base substitutions.

However, we observed a small enrichment of potential driver mutations in alcoholic liver ASCs (Fig. 3c; Table 1), although the number of mutations was low. Only one in three healthy liver ASCs acquired a nonsynonymous base substitution in a COSMIC cancer census gene. In alcoholic liver ASCs, on the other hand, we observed a total of seven nonsynonymous base substitutions in these cancer genes across eight ASCs. Two alcoholic liver ASCs even acquired multiple nonsynonymous hits in cancer genes (Fig. 3c; Table 1), while only an estimated four nonsynonymous base substitutions in cancer genes is sufficient to drive the development of liver cancer^16^. Consistent with this idea, we identified four nonsynonymous base substitutions in cancer genes in the MRCA of the HCC (Fig. 3c; Table 1). The modest increase in nonsynonymous base substitutions in cancer genes observed in alcoholic liver ASCs suggests that alcohol may cause clonal outgrowth of cells with putative oncogenic mutations, similar to alcohol-exposed esophagus^36^ and HCV-induced liver cirrhosis^21^.

**Table 1.** Somatic missense and nonsense base substitutions in cancer genes observed in healthy liver ASCs, alcoholic liver ASCs, and the MRCA of an HCC. *q* values (likelihood ratio test, FDR correction) indicate significant enrichment of nonsynonymous base substitutions within genes.

| Gene | Entrez<br>gene ID | Sample type | Sample(s) | Mutation type | *q*<br>(missense) | *q*<br>(nonsense) |
| --- | --- | --- | --- | --- | --- | --- |
| *ATP1A1* | 476 | Healthy liver | 15-a | missense | 1.00 | 1.00 |
| *PAX7* | 5081 | Healthy liver | 21-a | missense | 1.00 | 1.00 |
| *PREX2* | 80243 | Healthy liver | 21-b | missense | 1.00 | 1.00 |
| *PTPRK* | 5796 | Alcoholic liver | alc3-a, alc5-a | nonsense, nonsense | 1.00 | 0.02 |
| *ALK* | 238 | Alcoholic liver | alc3-b | missense | 1.00 | 1.00 |
| *CACNA1D* | 776 | Alcoholic liver | alc3-b | missense | 1.00 | 1.00 |
| *ZNF331* | 55422 | Alcoholic liver | alc4-b | missense | 1.00 | 1.00 |
| *CUX1* | 1523 | Alcoholic liver | alc5-a | missense | 1.00 | 1.00 |
| *TERT* | 7015 | Alcoholic liver | alc5-a | missense | 1.00 | 1.00 |
| *CD274* | 29126 | HCC | HCC-trunk | missense | 1.00 | 1.00 |
| *CIITA* | 4261 | HCC | HCC-trunk | missense | 1.00 | 1.00 |
| *KLF4* | 9314 | HCC | HCC-trunk | missense | 1.00 | 1.00 |
| *MUC1* | 4582 | HCC | HCC-trunk | missense | 1.00 | 1.00 |

Notably, we found that the cancer gene *PTPRK* was hit by a heterozygous nonsense base substitution in two alcoholic liver ASCs of independent patients, which is significantly more than expected based on the background mutation rate adjusted for the mutational profile (Table 1; likelihood ratio test, FDR correction; *q* = 0.02)^16^. None of the healthy liver ASCs acquired a nonsynonymous base substitution in *PTPRK*, nor did we identify nonsynonymous base substitutions in this gene in healthy ASCs from small intestine or colon^24^. RNA-sequencing of the organoids revealed that the heterozygous nonsense base substitutions in *PTPRK* resulted in a significantly reduced expression (Fig. 4; *P* < 0.05, negative binomial test), indicating a gene dosage effect due to nonsense-mediated decay of the mutated allele. *PTPRK* can modulate EGF-signaling through dephosphorylation of tyrosine residues of the EGFR^40^. Western blot analysis showed that alcoholic *PTPRK*^WT/*^ cells had increased pERK levels in the absence of EGF, indicating that the *PTPRK* mutations indeed disturb EGF-signaling (Fig. 4).

## DISCUSSION

In this study, we aimed to identify the mutational consequences of chronic alcohol intake in cirrhotic livers, prior to the development of liver cancer. In contrast to previous studies^12–15,37^, we did not observe specific mutational signatures associated with alcohol consumption in stem cells from non-cancerous cirrhotic livers. This observation can indicate that chronic alcohol use may only impact directly on mutation accumulation after tumor initiation or it may reflect very effective negative selection of cells with DNA damage. Alternatively, it should be noted that tissue-specific liver ASCs may not be the direct cell-of-origin for HCC. Nevertheless, as the cancer-initiating cells are exposed to the same mutagenic damage as the liver ASCs (and show the same mutational signatures), our results suggest that alcohol-induced cancer risk is not caused by altered mutagenesis.

We propose that chronic alcohol consumption creates an inflamed, cirrhotic liver tissue environment, which in turn provides a fertile ground for cells with specific oncogenic mutations to clonally expand. Chronic damage to the liver due to chronic alcohol consumption causes apoptosis and necrosis of various cells in the liver, such as the hepatocytes, leading to liver inflammation^41^. As a consequence, tissue-specific ASCs, which are normally quiescent, will proliferate to aid in the regeneration of the damaged liver^42^. Oncogenic mutations that have accumulated randomly in ASCs through normal mutational processes could provide a proliferative advantage under these inflamed conditions at the expense of ‘normal’ ASCs that do not carry such mutations, while there is too little proliferation under normal conditions for ASCs with oncogenic mutations to outcompete normal ASCs. In the damaged liver, the enhanced proliferation of such potential cancer progenitor cells could subsequently result in an increasing number of mutations that drive tumorigenesis further, as long as inflammation persists.

An illustrative example of this potentially altered selection process in the current study is the significant enrichment of nonsense base substitutions in EGFR phosphatase *PTPRK* in alcoholic liver ASCs. Reduced expression of *PTPRK* has been reported to cause enhanced EGF-signaling and ultimately increases cellular proliferation^40,43,44^. Single-nucleotide polymorphisms in *EGF* that prolong the half-life of EGF increase the risk of developing HCC through continued EGF-signaling^17,45,46^ and 22% of HCCs carry mutations in genes in the EGF pathway^47^. Reduced EGF-signaling, on the other hand, significantly decreases tumor formation in cirrhotic livers from rats^48^. These observations underscore the importance of (disturbed) EGF-signaling in HCC development. Nonsense mutations in *PTPRK* may thus contribute to the development of HCC by changing EGF-signaling as well, although further research should be conducted to identify the significance of our findings in liver cancer.

The liver is not the only organ in which inflammation is suggested to contribute to carcinogenesis. Inflammatory diseases, such as inflammatory bowel disease and pancreatitis, increase the risk of developing cancer in various tissues^49^. However, it was believed that this increased risk was at least partially due to a direct induction of mutations in the genome^50^. The results presented here indicate that alteration of selective mechanisms induced by inflammation could be more directly involved in the development of cancer. For liver, reversal of the inflammatory phenotype that precedes cancer might aid in reducing cancer risk in patients with cirrhotic liver disease due to chronic alcohol use.

## METHODS

### Human tissue material

All human tissue biopsies were obtained in the Erasmus MC - University Medical Center Rotterdam. Liver biopsies from healthy liver donors and patients with alcoholic cirrhosis were obtained during liver transplantation procedures. All patients were negative for viral infection and metabolic diseases. The biopsies were collected in cold organ preservation fluid (Belzer UW Cold Storage Solution, Bridge to Life, London, UK) and transported and stored at 4°C until use. The liver and tumor biopsies from the hepatocellular carcinoma patient were collected from a resected specimen and stored at -80°C until use. The acquisition of these liver and tumor biopsies for research purposes was approved by the Medical Ethical Committee of the Erasmus Medical Center (MEC-2014-060 and MEC-2013-143). Informed consent was provided by all patients involved.

The biopsies of the HCC were cut into 6µm sections. Subsequently, the tumor percentage of both ends of each biopsy was determined using HE staining (Supplementary Fig. 6). The tumor percentage of the biopsies was determined by averaging both values. The remaining slices were used for long-term storage at -80°C or for DNA isolation.

### Generation of clonal liver organoid cultures from human liver biopsies

Organoid cultures from healthy and alcoholic liver tissue material were derived as previously described^51,52^. After 2 - 3 days, organoids started to appear in the BME. The cultures were maintained for approximately 2 weeks after isolation, to enrich for ASCs. Subsequently, clonal organoid cultures were generated from these organoid cultures as described previously^25^. The organoid cultures were expanded until there was material for WGS.

### Whole-genome sequencing and read alignment

DNA was isolated from all organoid cultures, blood samples, and tissue biopsies using the Qiasymphony (Qiagen). Whole-genome sequencing libraries were generated from 200 ng of genomic DNA according to standard Illumina protocols. The organoid cultures and control samples were sequenced paired-end (2 × 150bp) to a depth of at least 30X coverage using the Illumina HiSeq Xten. The HCC biopsies were sequenced paired-end (2 × 150bp) to a depth of at least 60X coverage using the Illumina HiSeq Xten. A 60X depth was required to identify the somatic base substitutions in the tumor cells, as the biopsies contain ∼50% healthy cells (Supplementary Fig. 6). Whole-genome sequencing was performed at the Hartwig Medical Foundation in Amsterdam, the Netherlands. The sequence reads were mapped to the human reference genome GRCh37 using the Burrows–Wheeler Aligner (BWA) tool v0.7.5a^53^ (settings -t, 4, -c, 100, -M).

### Copy number alteration calling and filtering

For the healthy and alcoholic samples without HCC, CNA catalogs were obtained and filtered by using FreeC v2.7^25,54^. Calls were excluded if the mapping quality of the split reads was 0 on either sides of the split read. BED-file of blacklist positions is available upon request. For the HCC biopsies, structural variants were called using Manta v.1.1.0^55^ with standard settings. We only considered structural variations of at least 150 base pairs in autosomal the genome with a manta filter ‘PASS’. Subsequently, the mutation catalogs of all five biopsies were intersected with a window of 500 bp to obtain the trunk CNAs using bedtools^56^.

All CNA calls were inspected manually in the Integrative Genomics Viewer (IGV) to exclude false-positives with no change in read-depth. The breakpoints were identified manually in IGV. Finally, the number of genes within the deletions was obtained from http://genome.ucsc.edu/.

### Genome-wide copy number profiles

Genome-wide copy number profiles of the ASCs were estimated by using the output of the FreeC calls obtained in section ‘Copy number alteration calling and filtering’ prior to filtering. Subsequently, we calculated the mean copy number across 500,000 bp bins. Copy number of ≥ 2.8 was considered a gain and copy number of ≤ 1.2 a loss. Genome-wide copy number profiles of the HCC biopsies and the adjacent liver biopsy were obtained in a similar manner.

### Base substitution calling and filtering

For the organoid cultures, base substitution catalogs were obtained by filtering GATK v3.4-46^57^ variant calls as previously described^25^, with additional removal of variants with a sample-specific genotype quality < 10 in the control sample, and positions with a sample-specific genotype quality < 99 in the organoid clone sample. The callable regions, used to define regions with high confidence base substitutions, were obtained by using the GATK CallableLoci tool v3.4.46^58^ as previously described^25^. BED-file of blacklist positions is available upon request. All organoids showed a peak at a base substitution VAF of 0.5, confirming that the organoid samples are clonal (Supplementary Fig. 7). Publicly available variant call format (VCF) files and surveyed bed files of healthy liver ASCs were downloaded from donors 14 - 18 from https://wgs11.op.umcutrecht.nl/mutational_patterns_ASCs/ to allow the comparison between healthy and alcoholic liver ASCs.

For the HCC biopsies, base substitutions were called by using Strelka v1.0.14 with settings ‘SkipDepthFilters = 0’, ‘maxInputDepth = 250’, ‘depthFilterMultiple = 3.0’, ‘snvMaxFilteredBasecallFrac = 0.4’, ‘snvMaxSpanningDeletionFrac = 0.75’, ‘indelMaxRefRepeat = 1000’, ‘indelMaxWindowFilteredBasecallFrac = 0.3’, ‘indelMaxIntHpolLength = 14’, ‘ssnvPrior = 0.000001’, ‘sindelPrior = 0.000001’, ‘ssnvNoise = 0.0000005’, ‘sindelNoise = 0.000001’, ‘ssnvNoiseStrandBiasFrac = 0.5’, ‘minTier1Mapq = 20’, ‘minTier2Mapq = 5’, ‘ssnvQuality_LowerBound = 10’, ‘sindelQuality_LowerBound = 10’, ‘isWriteRealignedBam = 0’, and ‘binSize = 25000000’. We only considered variations with a filter ‘PASS’. Subsequently, the mutation catalogs of all five biopsies were intersected to obtain the trunk mutations using bedtools^56^. We only considered base substitutions on the autosomal genome that did not overlap with an indel call. Positions that were detected at least 5 times in 1,762 Dutch individuals were removed from these catalogs using the Hartwig Medical Foundation Pool of Normals (HMF-PON) version 2 (available upon request), to exclude Dutch germline variations. Only 138 base substitutions are found in four out of five biopsies, whereas we detect 7,203 base substitutions in all five biopsies (Supplementary Fig. 4), indicating that the majority of the trunk mutations were identified successfully.

To exclude that the observed similarities/differences in base substitution load and type are a consequence of the differences between the filtering pipelines, we also applied the filtering steps of the HCC samples to the base substitutions in the alcohol liver ASCs. We observe no obvious differences in base substitution load or type between the alcoholic liver samples using both filtering pipelines (Supplementary Fig. 8).

### Tumor adjusted allele frequencies

The VAFs of the shared base substitutions (the trunk mutations) were calculated for each biopsy. Subsequently, we calculated the tumor-adjusted variant allele frequency (TAF) per biopsy, in which the VAF is divided by the tumor-fraction. Chromosome 1 and chromosome 8 were excluded from these analyses, as these chromosomes deviate from a copy number of two in the majority of the biopsies (Supplementary Fig. 4). Most biopsies showed a peak around a TAF of 0.5 (Supplementary Fig. 9), confirming that these base substitutions are clonal in each sample and that the biopsies share a recent common ancestor.

### Indel calling and filtering

WGS data of previously published samples was obtained from EGAD00001001900. Indels were called using GATK v3.4-46^57^. We only considered indels that were callable/surveyed on autosomal chromosomes with one alternative allele and a GATK filter ‘PASS’. To remove false positive calls, indels with a GATK quality score < 250 and/or with a mapping quality < 60 were excluded. Additionally, only indels with a coverage of at least 20X and a GATK sample-specific quality score of at least 99 in both control and organoid clone sample were considered. Subsequently, variants with a cosmic and/or a dbSNP id (dbSNP v137.b3730) and indels that were found in three unrelated control samples (BED-file available upon request) were excluded. To obtain a catalog of somatic indels, we excluded indels with any evidence in the reference sample, and that were located within 100 base pairs of an indel that was called in the reference sample. Finally, we only considered variants with a VAF of ≥ 0.3 in the organoid clone sample.

### (Tandem) base substitution and indel rate in liver ASCs

The number of base substitutions in the genomes of liver ASCs was obtained from the VCF files and extrapolated to the non-N autosomal genome (2,682,655,440 bp) of GRCh37 using the callable/surveyed genome size obtained in section ‘Base substitution calling and filtering’. To identify whether the number of somatic base substitutions acquired in the genomes of liver ASCs are correlated with the age of the donor, we fitted a linear mixed-effects regression model with the donor as a random effect in this model using the nlme R package, as described previously^24^. Two-tailed *t*-tests were performed to determine whether the correlation between age and number of mutations was significant. The accumulation of base substitutions did not correlate significantly with age in the alcoholic liver ASCs (∼38.6 somatic base substitutions per year; 95% CI: -51.8 - 128.9; two-tailed *t*-test, linear mixed model, non-significant), most likely due to the fact that the age-range is much smaller in these donors. Therefore, we obtained the 95% CI of the healthy liver ASCs from the output of the linear mixed-effects regression model and determined whether the number of somatic base substitutions acquired in the genomes of the alcoholic liver ASCs are within this 95% CI.

To identify tandem base substitutions, we extracted base substitutions that were called on two consecutive bases in the GRCh37 human reference genome from the VCF files. Similar to single base substitutions, we extrapolated this number to the non-N autosomal genome and determined whether the number of tandem base substitutions was correlated with the age of the donor using a linear mixed effects regression model. As the number of tandem base substitutions did not significantly correlate with age in the alcoholic liver ASCs (∼0.04 tandem base substitutions per year; 95% CI: -1.13 - 1.21; two-tailed *t*-test, linear mixed model, non-significant), we determined whether these tandem base substitution numbers are within the 95% CI of the healthy liver ASCs.

Similar to base substitutions and tandem base substitutions, the number of indels was extracted from the filtered VCF files and extrapolated to the non-N autosomal genome. Using a linear mixed effects regression model with the donor as random effect, we assessed whether the number of indels was correlated with the age of the donor. Two-tailed *t*-test were performed to determine whether the correlation was significant for both alcoholic liver ASCs and healthy liver ASCs.

### Mutational pattern analysis

Mutation types of the base substitutions were extracted from the VCF files and the mutational profiles were generated by retrieving the sequence context of each base substitution. For the healthy and alcoholic liver ASCs, we calculated an ‘average’ mutational profile. Pairwise cosine similarities of these average mutational profiles and of the mutational profile of the trunk mutations of the HCC were calculated, to identify the similarity between these profiles.

We reconstructed the mutational profiles of the average mutational profiles and the trunk mutations using the 60 known SBS signatures (Supplementary Fig. 3; https://cancer.sanger.ac.uk/cosmic/signatures v3). A bootstrap resampling method similar as described in Zou *et al*., ^35^ was used to generate 120,000 (8 × 15 x 1,000) replicas of the mutational profiles of the healthy and alcoholic liver ASCs. Subsequently, 8 or 15 (for healthy and alcoholic liver ASCs respectively) replicas were randomly selected and an average mutational profile was calculated. This was repeated 10,000 times, to obtain 10,000 average mutational profiles of the replicas for both healthy and alcoholic livers. These average mutational profiles of the replicas were reconstructed using the 60 known SBS mutational signatures and the Euclidean distance to the original signature contribution was calculated for each reconstructed average mutational profile. Next, the distance at which *P* = 0.01 was determined for both healthy and alcoholic liver ASCs (*d*_healthy=0.01_ and *d*_alcoholic=0.01_, respectively). The Euclidean distance (*d*) between the original signature contributions of healthy and alcoholic liver ASCs was considered significant when *d* was larger than *d*_healthy=0.01_ and *d*_alcoholic=0.01_.

### Genomic distribution of somatic base substitutions

The promoter, enhancer, and open chromatin regions of hg19 were obtained from Ensembl using biomaRt^59,60^ and the genic regions of hg19 were loaded from UCSC Known Genes tables as TxDb object^61^. To determine whether the somatic base substitutions are non-randomly distributed, we tested for enrichment and depletion of base substitutions in these regions with a one-sided Binomial test, corrected for the callable/surveyed regions per sample, similar as described in ^24^. For the HCC trunk mutations, the callable regions were obtained by defining callable loci per biopsy using the GATK CallableLoci tool v3.4.46^58^ (optional parameters ‘minBaseQuality 10’, ‘minMappingQuality 10’, ‘maxFractionOfReadsWithLowMAPQ 20’, and ‘minDepth 15’). Subsequently, these files were intersected to obtain the regions that are callable in all biopsies. 96.79% of the non-N autosomal genome was callable in all six biopsies. Two-sided poisson tests were done to estimate significant differences in depletion/enrichment in all genomic regions between the healthy liver ASCs, the alcoholic liver ASCs, and the trunk mutations of the HCC. Differences were considered significant when *q* < 0.05 (Benjamini-Hochberg FDR multiple-testing correction). All mutational pattern analyses were performed using the MutationalPatterns R package^62^.

To obtain a generic genome-wide profile of H3K36me3, we downloaded and merged 40 available H3K36me3 ChIP-Seq datasets from UCSC, and determined the median H3K36me3 values in regions that show H3K36me3 enrichment in at least 2 of the datasets. H3K36me3 peaks were subsequently called using bdgpeakcall function of MACS2 (broad peaks)^63^. The amount of base substitutions that overlap with these peaks was calculated for all base substitutions acquired in liver ASCs. These analyses were repeated for T:A > C:G mutations specifically. Wilcoxon-rank tests were performed to estimate significant differences in the relative amount of base pair substitutions in H3K36me3 regions between alcoholic and healthy liver ASCs. Differences with *P* < 0.05 were considered significant.

### d*N*/d*S* and identification of nonsynonymous base substitutions in cancer genes

d*N*/d*S* ratios were computed using the *dNdScv* R package^16^. The output of the *dNdScv* package was used to identify missense, nonsense, and splice site base substitutions in cosmic cancer genes. For this analysis, we considered all 409 ‘tier 1’ cancer genes (genes with sufficient evidence of being a cancer driver). The list of cosmic cancer genes was obtained from https://cancer.sanger.ac.uk/cosmic/census.

### RNA sequencing

Organoid cultures of three healthy donors (18-c, 21-b, and 22-a) and three alcoholic organoids, of which 2 with a nonsense base substitution in *PTPRK* (alc3-a and alc5-a) and one without any base substitutions in *PTPRK* (alc-3b), were cultured for 1 day either in presence or absence of hEGF in the culture medium. Subsequently, cells were collected in Trizol. Total RNA was isolated using the QiaSymphony SP with the QiaSymphony RNA kit (Qiagen, 931636). mRNA sequencing libraries were generated from 50 ng total RNA using the Illumina Neoprep TruSeq stranded mRNA library prep kit (Illumina, NP-202-1001). RNA libraries were sequenced paired-end (2 × 75 bp) on the Nextseq500 to > 20 million reads per sample at the Utrecht Sequencing facility.

RNA sequencing reads were mapped to the human reference genome GRCh37 with STAR v.2.4.2a^64^. The BAM-files were indexed using Sambamba v0.5.8 Subsequently, reads were counted using HTSeq-count 0.6.1 and read counts were normalized using DESeq v1.18.0. DESeq nbinomTest was used to test for differential expression of *PTPRK* between the organoids with a nonsense *PTPRK* base substitution and the other organoids.

### Western blot

Simultaneous to the collection of samples for RNA isolation described above, we also obtained protein samples for western blot in Laemmli buffer. 20 ug of protein was run on a 10% SDS page gel and blocked for 1 hour using 5% ELK in TBS-T after transfer to a nitrocellulose membrane. Subsequently, the membrane was incubated overnight with primary antibody (pERK AB50011, abcam; ERK AB17942, abcam; Actin A2066, sigma-aldrich) and for 1 hour at room temperature with secondary antibody. We visualized the proteins with the Amersham ECL Western blotting analysis system (GE Healthcare, RPN2109).

## Data availability

The whole-genome sequencing and RNA sequencing data generated during the current study are available at EGA (https://www.ebi.ac.uk/ega/home) under accession number EGAS00001002983. Filtered VCF-files, metadata, BED-files with callable regions, and RNA-Seq counts generated during the current study are available at Zenodo under DOI 10.5281/zenodo.3295513 (https://doi.org/10.5281/zenodo.3295513). Data analysis scripts used during the current study are available at https://github.com/UMCUGenetics/Liverdisease, https://github.com/UMCUGenetics/IAP and https://github.com/hartwigmedical.

## ACKNOWLEDGEMENTS

The authors would like to thank the Utrecht Sequencing Facility and the UBEC for sequencing and for input on the bioinformatic analyses, respectively. The UBEC is subsidized by the University Medical Center Utrecht and the Utrecht Sequencing Facility is subsidized by the University Medical Center Utrecht, Hubrecht Institute, and Utrecht University. This study was financially supported by the research program InnoSysTox (project number 114027003), by the Netherlands Organisation for Health Research and Development (ZonMw), by the Dutch Cancer Society (project number 10496) and is part of the Oncode Institute, which is partly financed by the Dutch Cancer Society and was funded by the gravitation program CancerGenomiCs.nl from the Netherlands Organisation for Scientific Research (NWO). We thank the Hartwig Medical Foundation (Amsterdam, The Netherlands) for generating, analyzing and providing access to reference whole genome sequencing data of the Netherlands population.

## AUTHOR CONTRIBUTIONS

R.L., J.J., J.I., M.D., and M.V. collected liver biopsies. M.J., E.K., and N.B. performed organoid culturing. N.B. isolated the RNA and protein, prepared RNA-seq libraries of the organoid cultures and performed Western blot. M.J., M.L., B.R., R.J., and S.B. performed bioinformatic analyses. M.J., E.K., M.V., R.B., L.L., and E.C. were involved in the conceptual design of this study. M.J. and E.C. wrote the manuscript. All authors provided textual comments and have approved the manuscript. R.B., L.L., and E.C. supervised this study.

## AUTHOR INFORMATION

### Corresponding authors

Correspondence to Edwin Cuppen.

## REFERENCES

1. Stewart, B. W. & Wild, C. P. World Cancer Report 2014. (2014).

2. Boffetta, P., Hashibe, M., La Vecchia, C., Zatonski, W. & Rehm, J. The burden of cancer attributable to alcohol drinking. International Journal of Cancer 119, 884–887 (2006).

3. World Health Organization. Global Status Report on Alcohol and Health. (World Health Organization, 2014).

4. Mizumoto, A. et al. Molecular Mechanisms of Acetaldehyde-Mediated Carcinogenesis in Squamous Epithelium. Int. J. Mol. Sci. 18, (2017).

5. Obe, G. & Ristow, H. Mutagenic, cancerogenic and teratogenic effects of alcohol. Mutat. Res. 65, 229–259 (1979).

6. Helander, A. & Lindahl-Kiessling, K. Increased frequency of acetaldehyde-induced sister-chromatic exchanges in human lymphocytes treated with an aldehyde dehydrogenase inhibitor. Mutat. Res. Lett. 264, 103–107 (1991).

7. Matsuda, T., Kawanishi, M., Matsui, S., Yagi, T. & Takebe, H. Specific tandem GG to TT base substitutions induced by acetaldehyde are due to intra-strand crosslinks between adjacent guanine bases. Nucleic Acids Res. 26, 1769–1774 (1998).

8. Tamura, M., Ito, H., Matsui, H. & Hyodo, I. Acetaldehyde is an oxidative stressor for gastric epithelial cells. J. Clin. Biochem. Nutr. 55, 26–31 (2014).

9. Novitskiy, G., Traore, K., Wang, L., Trush, M. A. & Mezey, E. Effects of ethanol and acetaldehyde on reactive oxygen species production in rat hepatic stellate cells. Alcohol. Clin. Exp. Res. 30, 1429–1435 (2006).

10. Grollman, A. P. & Moriya, M. Mutagenesis by 8-oxoguanine: an enemy within. Trends Genet. 9, 246–249 (1993).

11. van Loon, B., Markkanen, E. & Hübscher, U. Oxygen as a friend and enemy: How to combat the mutational potential of 8-oxo-guanine. DNA Repair 9, 604–616 (2010).

12. Chang, J. et al. Genomic analysis of oesophageal squamous-cell carcinoma identifies alcohol drinking-related mutation signature and genomic alterations. Nat. Commun. 8, 15290 (2017).

13. Schulze, K. et al. Exome sequencing of hepatocellular carcinomas identifies new mutational signatures and potential therapeutic targets. Nat. Genet. 47, 505–511 (2015).

14. Fujimoto, A. et al. Whole-genome mutational landscape and characterization of noncoding and structural mutations in liver cancer. Nat. Genet. 48, 500–509 (2016).

15. Letouzé, E. et al. Mutational signatures reveal the dynamic interplay of risk factors and cellular processes during liver tumorigenesis. Nat. Commun. 8, 1315 (2017).

16. Martincorena, I. et al. Universal Patterns of Selection in Cancer and Somatic Tissues. Cell 171, 1029–1041.e21 (2017).

17. Hernandez–Gea, V., Toffanin, S., Friedman, S. L. & Llovet, J. M. Role of the Microenvironment in the Pathogenesis and Treatment of Hepatocellular Carcinoma. Gastroenterology 144, 512–527 (2013).

18. Zhu, L. et al. Multi-organ Mapping of Cancer Risk. Cell 166, 1132–1146.e7 (2016).

19. Seitz, H. K. & Stickel, F. Molecular mechanisms of alcohol-mediated carcinogenesis. Nat. Rev. Cancer 7, 599–612 (2007).

20. Desai, A., Sandhu, S., Lai, J.-P. & Sandhu, D. S. Hepatocellular carcinoma in non-cirrhotic liver: A comprehensive review. World Journal of Hepatology 11, 1–18 (2019).

21. Zhu, M. et al. Somatic Mutations Increase Hepatic Clonal Fitness and Regeneration in Chronic Liver Disease. Cell (2019). doi:10.1016/j.cell.2019.03.026

22. Khosla, R. et al. EpCAM+ Liver Cancer Stem-Like Cells Exhibiting Autocrine Wnt Signaling Potentially Originate in Cirrhotic Patients. Stem Cells Transl. Med. 6, 807–818 (2017).

23. Chen, D. et al. Epithelial to mesenchymal transition is involved in ethanol promoted hepatocellular carcinoma cells metastasis and stemness. Mol. Carcinog. 57, 1358–1370 (2018).

24. Blokzijl, F. et al. Tissue-specific mutation accumulation in human adult stem cells during life. Nature 538, 260–264 (2016).

25. Jager, M. et al. Measuring mutation accumulation in single human adult stem cells by whole-genome sequencing of organoid cultures. Nat. Protoc. 13, 59–78 (2018).

26. Barker, N. et al. Crypt stem cells as the cells-of-origin of intestinal cancer. Nature 457, 608–611 (2009).

27. Adams, P. D., Jasper, H. & Rudolph, K. L. Aging-Induced Stem Cell Mutations as Drivers for Disease and Cancer. Cell Stem Cell 16, 601–612 (2015).

28. Lee, J.-S. et al. A novel prognostic subtype of human hepatocellular carcinoma derived from hepatic progenitor cells. Nat. Med. 12, 410–416 (2006).

29. Tummala, K. S. et al. Hepatocellular Carcinomas Originate Predominantly from Hepatocytes and Benign Lesions from Hepatic Progenitor Cells. Cell Rep. 19, 584–600 (2017).

30. Holczbauer, Á. et al. Modeling pathogenesis of primary liver cancer in lineage-specific mouse cell types. Gastroenterology 145, 221–231 (2013).

31. Mu, X. et al. Hepatocellular carcinoma originates from hepatocytes and not from the progenitor/biliary compartment. Journal of Clinical Investigation 125, 3891–3903 (2015).

32. Alexandrov, L. B. et al. Signatures of mutational processes in human cancer. Nature 500, 415–421 (2013).

33. Nik-Zainal, S. et al. Landscape of somatic mutations in 560 breast cancer whole-genome sequences. Nature 534, 47–54 (2016).

34. Alexandrov, L. B. et al. The Repertoire of Mutational Signatures in Human Cancer. bioRxiv 322859 (2018). doi:10.1101/322859

35. Zou, X. et al. Validating the concept of mutational signatures with isogenic cell models. Nat. Commun. 9, 1744 (2018).

36. Yokoyama, A. et al. Age-related remodelling of oesophageal epithelia by mutated cancer drivers. Nature 565, 312–317 (2019).

37. Supek, F. & Lehner, B. Clustered Mutation Signatures Reveal that Error-Prone DNA Repair Targets Mutations to Active Genes. Cell 170, 534–547.e23 (2017).

38. Barski, A. et al. High-resolution profiling of histone methylations in the human genome. Cell 129, 823–837 (2007).

39. Bannister, A. J. et al. Spatial distribution of di- and trimethyl lysine 36 of histone H3 at active genes. J. Biol. Chem. 280, 17732–17736 (2005).

40. Xu, Y., Tan, L.-J., Grachtchouk, V., Voorhees, J. J. & Fisher, G. J. Receptor-type protein-tyrosine phosphatase-kappa regulates epidermal growth factor receptor function. J. Biol. Chem. 280, 42694–42700 (2005).

41. Luedde, T., Kaplowitz, N. & Schwabe, R. F. Cell death and cell death responses in liver disease: mechanisms and clinical relevance. Gastroenterology 147, 765–783.e4 (2014).

42. Lu, W.-Y. et al. Hepatic progenitor cells of biliary origin with liver repopulation capacity. Nat. Cell Biol. 17, 971–983 (2015).

43. Sun, P.-H., Ye, L., Mason, M. D. & Jiang, W. G. Protein tyrosine phosphatase kappa (PTPRK) is a negative regulator of adhesion and invasion of breast cancer cells, and associates with poor prognosis of breast cancer. J. Cancer Res. Clin. Oncol. 139, 1129–1139 (2013).

44. Flavell, J. R. et al. Down-regulation of the TGF-beta target gene, PTPRK, by the Epstein-Barr virus encoded EBNA1 contributes to the growth and survival of Hodgkin lymphoma cells. Blood 111, 292–301 (2008).

45. Zhong, J.-H. et al. Epidermal Growth Factor Gene Polymorphism and Risk of Hepatocellular Carcinoma: A Meta-Analysis. PLoS One 7, e32159 (2012).

46. Tanabe, K. K. et al. Epidermal growth factor gene functional polymorphism and the risk of hepatocellular carcinoma in patients with cirrhosis. JAMA 299, 53–60 (2008).

47. Sanchez-Vega, F. et al. Oncogenic Signaling Pathways in The Cancer Genome Atlas. Cell 173, 321–337.e10 (2018).

48. Schiffer, E. et al. Gefitinib, an EGFR inhibitor, prevents hepatocellular carcinoma development in the rat liver with cirrhosis. Hepatology 41, 307–314 (2005).

49. Mantovani, A., Allavena, P., Sica, A. & Balkwill, F. Cancer-related inflammation. Nature 454, 436–444 (2008).

50. Shimizu, T., Marusawa, H., Endo, Y. & Chiba, T. Inflammation-mediated genomic instability: roles of activation-induced cytidine deaminase in carcinogenesis. Cancer Sci. 103, 1201–1206 (2012).

51. Broutier, L. et al. Culture and establishment of self-renewing human and mouse adult liver and pancreas 3D organoids and their genetic manipulation. Nat. Protoc. 11, 1724–1743 (2016).

52. Huch, M. et al. Long-term culture of genome-stable bipotent stem cells from adult human liver. Cell 160, 299–312 (2015).

53. Li, H. & Durbin, R. Fast and accurate short read alignment with Burrows-Wheeler transform. Bioinformatics 25, 1754–1760 (2009).

54. Boeva, V. et al. Control-FREEC: a tool for assessing copy number and allelic content using next-generation sequencing data. Bioinformatics 28, 423–425 (2012).

55. Chen, X. et al. Manta: rapid detection of structural variants and indels for germline and cancer sequencing applications. Bioinformatics 32, 1220–1222 (2016).

56. Quinlan, A. R. BEDTools: The Swiss-Army Tool for Genome Feature Analysis. Curr. Protoc. Bioinformatics 47, 11.12.1–34 (2014).

57. McKenna, A. et al. The Genome Analysis Toolkit: a MapReduce framework for analyzing next-generation DNA sequencing data. Genome Res. 20, 1297–1303 (2010).

58. Van der Auwera, G. A. et al. From FastQ data to high confidence variant calls: the Genome Analysis Toolkit best practices pipeline. Curr. Protoc. Bioinformatics 43, 11.10.1–33 (2013).

59. Durinck, S. et al. BioMart and Bioconductor: a powerful link between biological databases and microarray data analysis. Bioinformatics 21, 3439–3440 (2005).

60. Durinck, S., Spellman, P. T., Birney, E. & Huber, W. Mapping identifiers for the integration of genomic datasets with the R/Bioconductor package biomaRt. Nat. Protoc. 4, 1184–1191 (2009).

61. Carlson, M. & Maintainer, B. P. TxDb.Hsapiens.UCSC.hg19.knownGene: Annotation package for TxDb object(s). (2015).

62. Blokzijl, F., Janssen, R., van Boxtel, R. & Cuppen, E. MutationalPatterns: comprehensive genome-wide analysis of mutational processes. Genome Med. 10, 33 (2018).

63. Zhang, Y. et al. Model-based analysis of ChIP-Seq (MACS). Genome Biol. 9, R137 (2008).

64. Dobin, A. et al. STAR: ultrafast universal RNA-seq aligner. Bioinformatics 29, 15–21 (2013).

