## Supplementary material for "Mutational impact of chronic alcohol use on stem cells in cirrhotic liver"

### Table of contents

#### Supplementary Figures

Supplementary Figure 1. Somatic mutations in genomes of healthy and alcoholic liver stem cells.

Supplementary Figure 2. Mutational profiles of the somatic base substitutions in the genomes of healthy and alcoholic liver ASCs.

Supplementary Figure 3. Reconstruction of mutational profiles with 60 COSMIC mutational signatures.

Supplementary Figure 4. Somatic mutations in biopsies of an alcohol-associated HCC.

Supplementary Figure 5. Boxplots depicting relative number of base substitutions and T:A>C:G mutations in H3K36Me3 enriched regions in healthy and alcoholic liver ASCs.

Supplementary Figure 6. HE-staining of slices of the ends of five HCC biopsies and of a healthy adjacent biopsy.

Supplementary Figure 7. Variant allele frequency (VAF) distributions of the somatic base substitutions acquired in the genomes of healthy and alcoholic liver ASCs before applying the VAF ≥ 0.3 filter.

Supplementary Figure 8. Comparison between the base substitution catalogs of the alcoholic liver ASCs obtained using two independent variant calling pipelines.

Supplementary Figure 9. Distribution of the variant allele frequencies of 6,232 somatic base substitutions that are shared by all five HCC biopsies, per biopsy, adjusted for the estimated tumor percentage per biopsy.

#### Supplementary Tables

Supplementary Table 1. Somatic base substitutions, tandem base substitutions, indels, and copy number alterations acquired in the genomes of healthy and alcoholic liver ASCs.

Supplementary Table 2. Somatic base substitutions identified in five biopsies of one alcohol-related HCC.

Supplementary Table 3. Somatic copy number alterations detected in a recent common ancestor of an alcohol-related HCC.

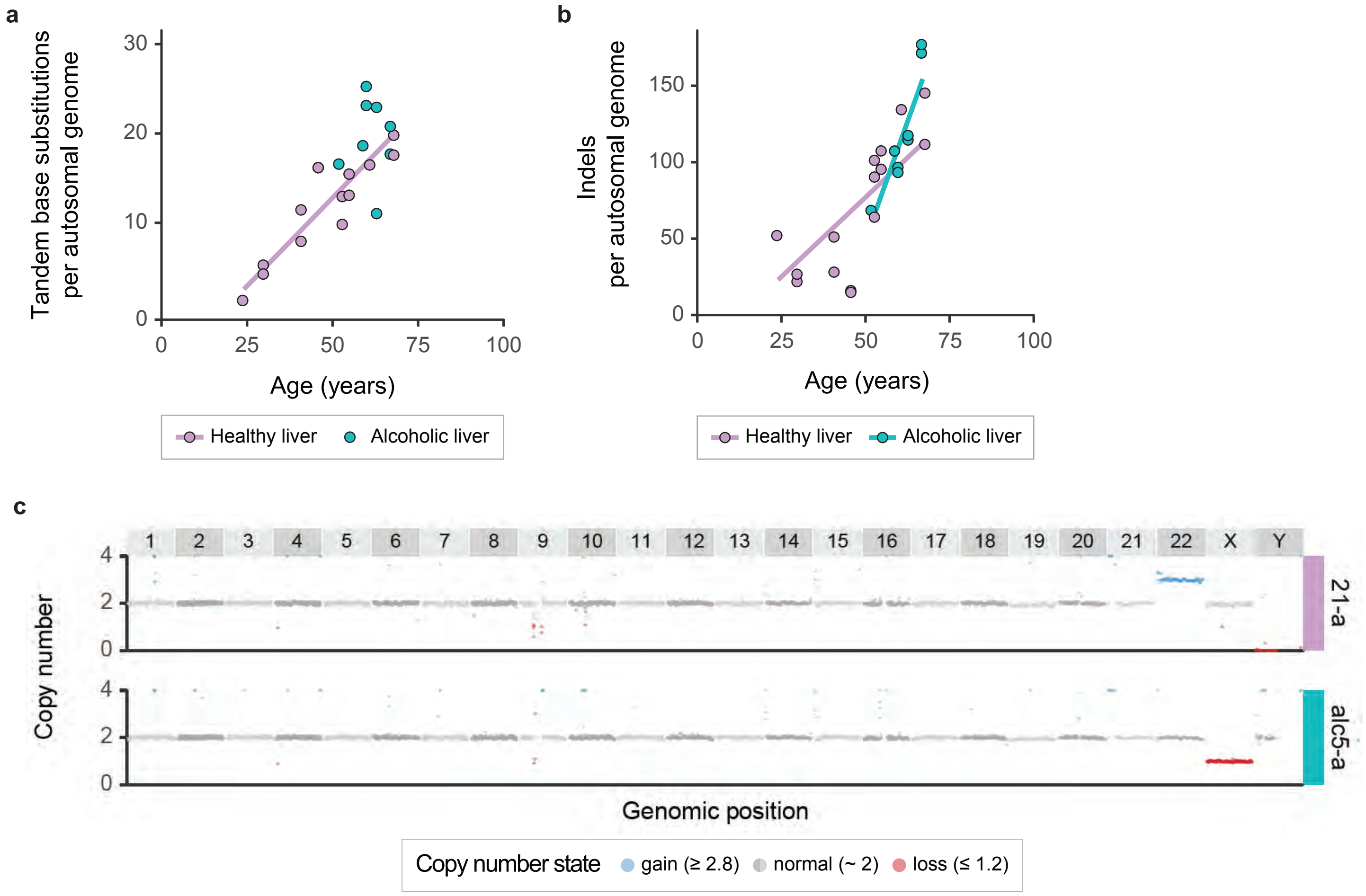


**Supplementary Figure 1.** Somatic mutations in genomes of healthy and alcoholic liver stem cells. Number of **a** tandem base substitutions and **b** indels in the autosomal genomes of 15 healthy and 8 alcoholic liver ASCs of 9 and 5 donors, respectively. Each stem cell is represented by a data point. A linear accumulation of tandem base substitutions with age was observed in the healthy liver (two-tailed *t*-test, linear mixed model, $P = 2.01 \times 10^{-4}$) and of indels with age in both healthy and alcoholic liver (two-tailed *t*-test, linear mixed model, $P = 1.79 \times 10^{-2}$ and $P = 2.35 \times 10^{-2}$, respectively), indicated by the purple and blue trendlines. Healthy liver ASCs acquire ~0.38 tandem base substitutions (95% CI: 0.25 - 0.51) and ~2.0 somatic indels (95% CI: 0.5 - 3.7) per year. Alcoholic liver ASCs acquire ~6.2 somatic indels (95% CI: 1.6 - 10.8) per year. All tandem base substitution numbers in the genomes of the alcoholic liver ASCs are within the 95% confidence interval of the healthy liver ASCs, except for alc4-b, which is slightly lower (11.3 tandem base substitutions at 63 years). **c** Genomic copy number profiles of ASC 21-a and alc5-a.

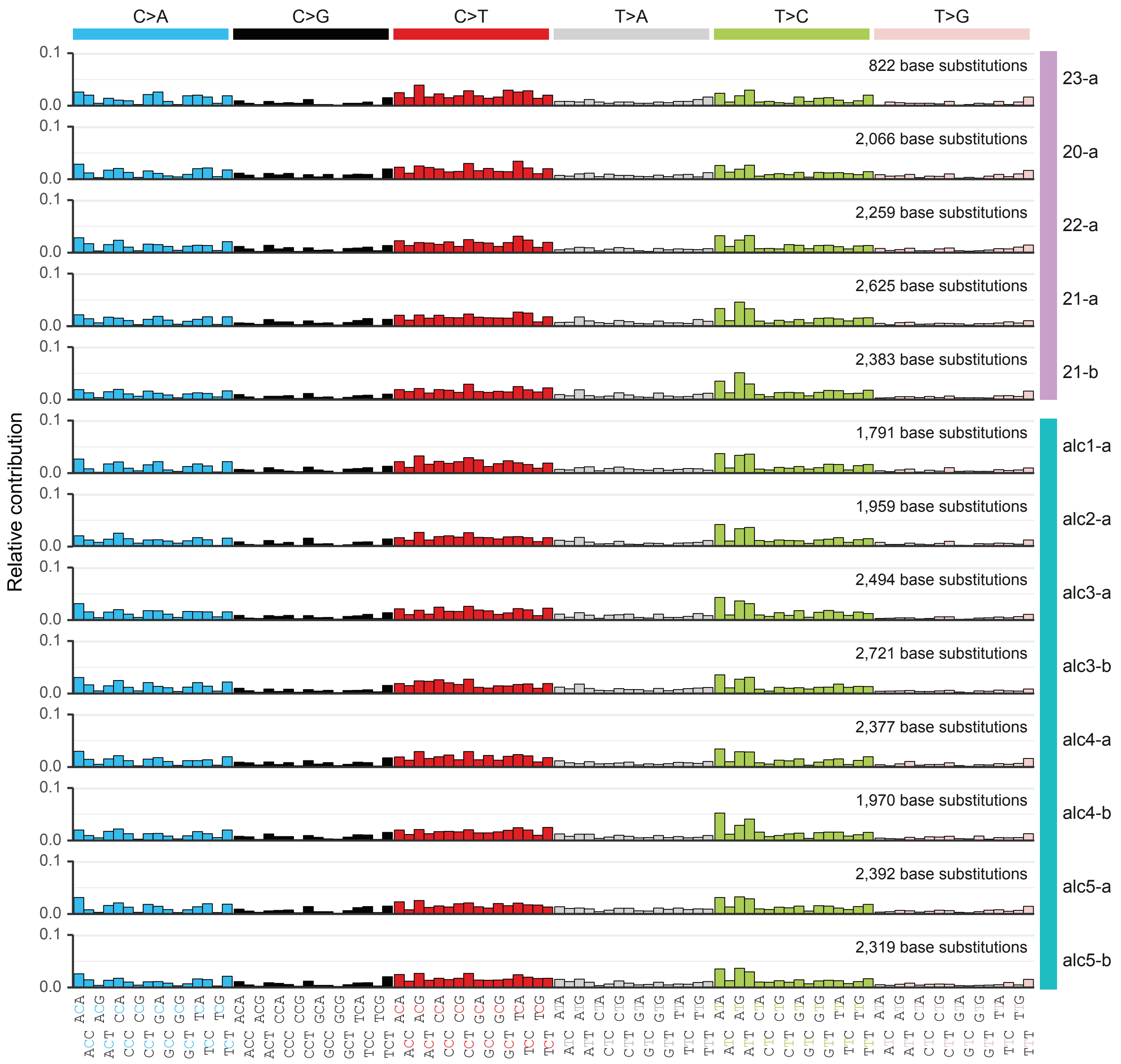


**Supplementary Figure 2.** Mutational profiles of the somatic base substitutions in the genomes of healthy and alcoholic liver ASCs. In this figure, we only displayed the mutational profiles of the new liver ASCs. Mutation spectra of the previously published healthy liver ASCs can be obtained from Extended Data Figure 5 of Blokzijl *et al*. [24].

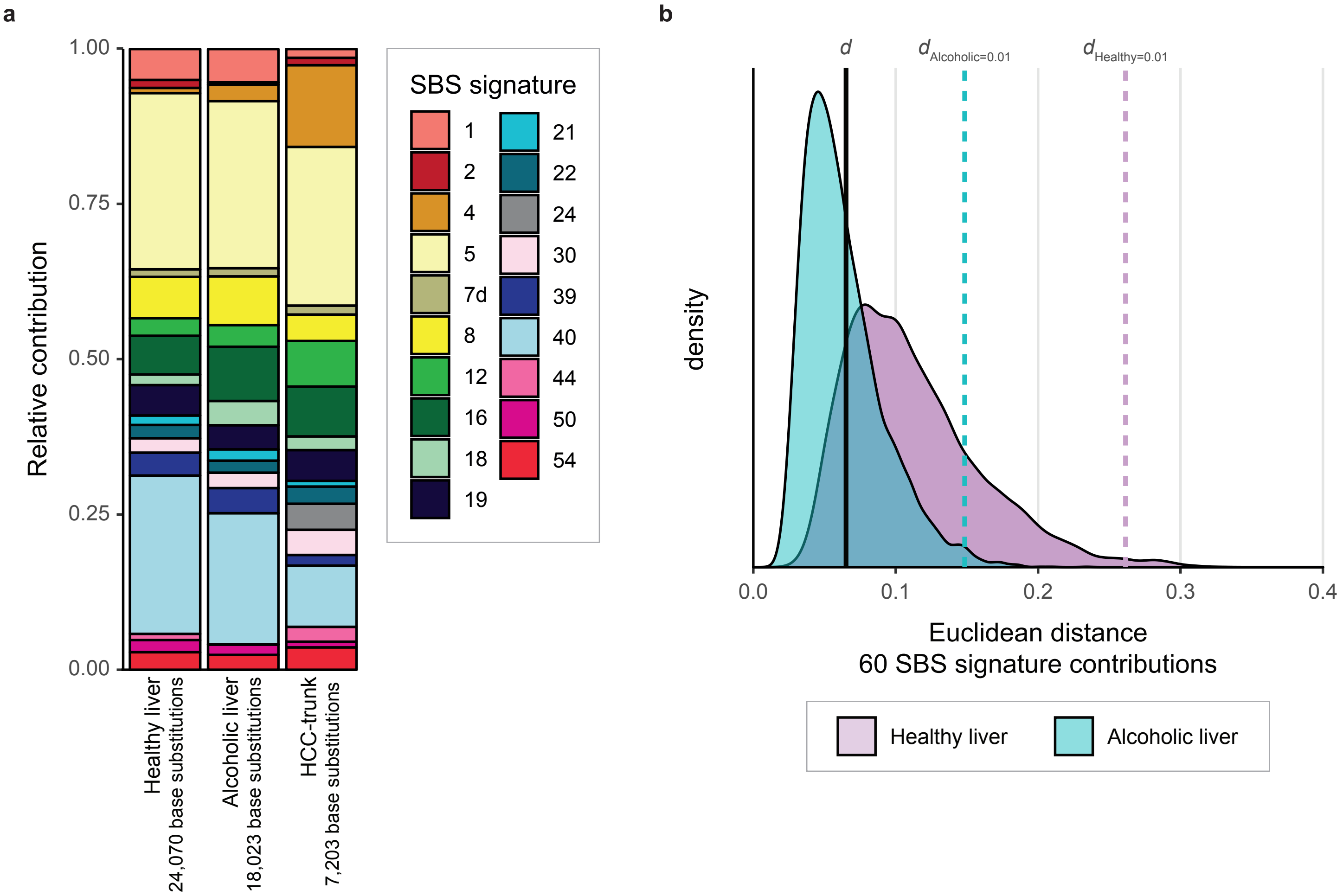


**Supplementary Figure 3.** Reconstruction of mutational profiles with 60 COSMIC mutational signatures. **a** Mean relative contribution of the 60 COSMIC SBS signatures (v3) to the mutational profiles of the somatic base substitutions of healthy and alcoholic liver ASCs and of the MRCA of an HCC. Only signatures with at least 100 mutations across the mutational profiles of the depicted sample types (mean mutational profiles of healthy and alcoholic liver ASCs, and HCC MRCA) are shown. **b** A bootstrap resampling method (see methods) was used to create a distribution of 10,000 signature contributions for both healthy and alcoholic ASCs. The Euclidean distance between each of these reconstructed signature contributions and the original signature contribution (depicted in **a**) is shown in purple for healthy and in blue for alcoholic liver ASCs. $d$ indicates the Euclidean distance between the original signature contributions of healthy and alcoholic liver ASC. The purple dashed line indicates the distance where $P$ = 0.01 for $d_{Healthy=0.01}$ and the blue dashed line indicates the distance where $P$ = 0.01 for $d_{Alcoholic=0.01}$. We considered the signature contribution to differ significantly between healthy and alcoholic liver ASCs, when $d$ is larger than $d_{Healthy=0.01}$ and $d_{Alcoholic=0.01}$.

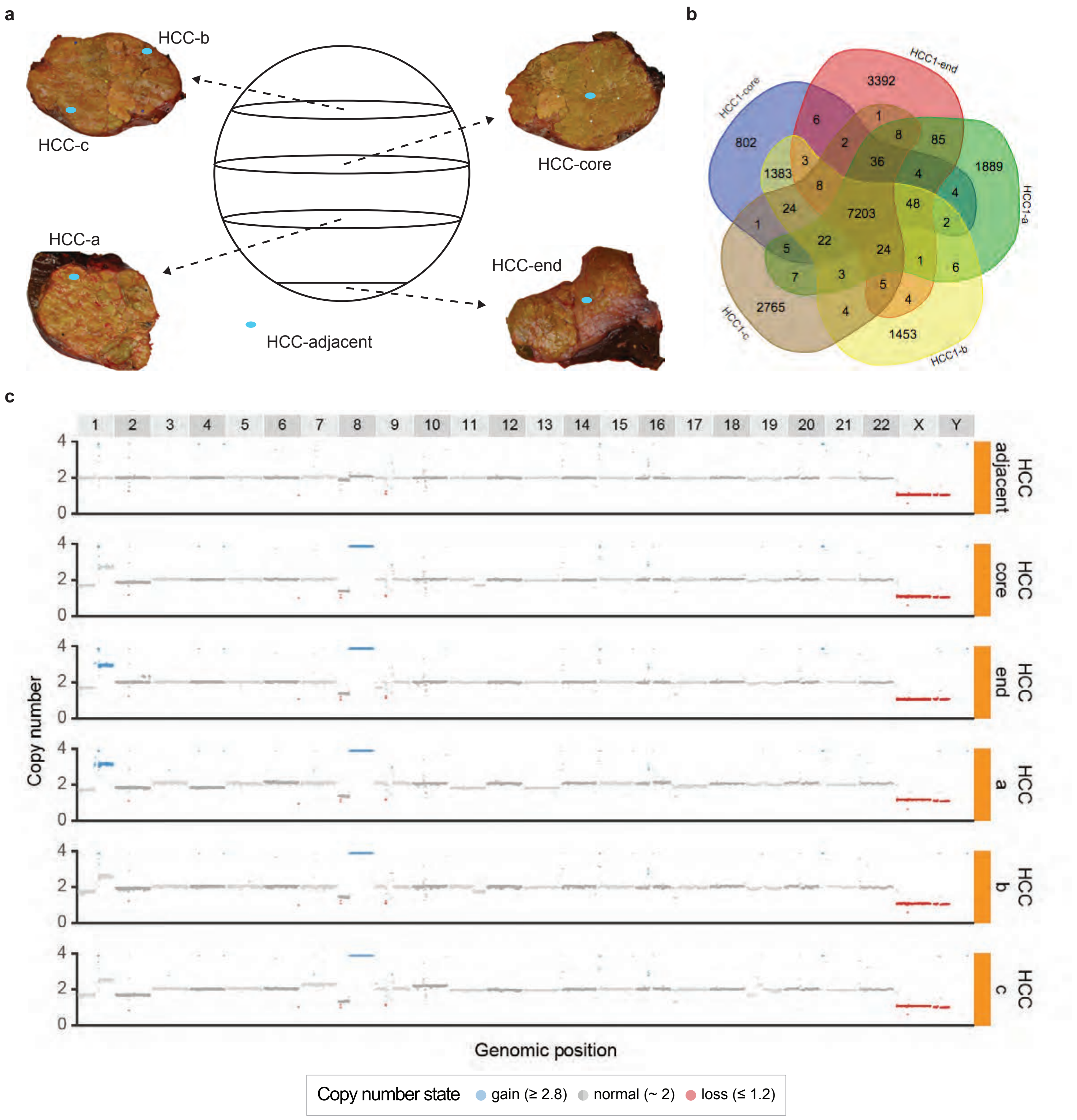


**Supplementary Figure 4.** Somatic mutations in biopsies of an alcohol-associated HCC. **a** Schematic depiction of the tumor (large circle) and the 4 slices of the tumor (ovals) of which biopsies were obtained. Biopsies are indicated by blue dots on the pictures of the HCC slices. **b** Venn diagram of somatic base substitutions identified in five biopsies of the alcohol-related HCC. Venn diagram was created using http://bioinformatics.psb.ugent.be/webtools/Venn/. **c** Genomic copy number profiles of all five HCC biopsies and of healthy adjacent tissue.

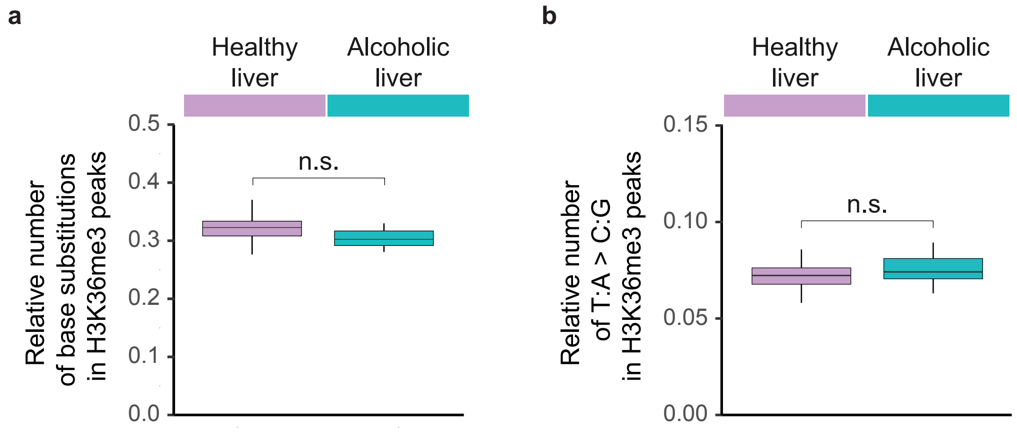


**Supplementary Figure 5.** Boxplots depicting relative number of **a** base substitutions and **b** T:A>C:G mutations in H3K36Me3 enriched regions in healthy and alcoholic liver ASCs. n.s. = non-significant.

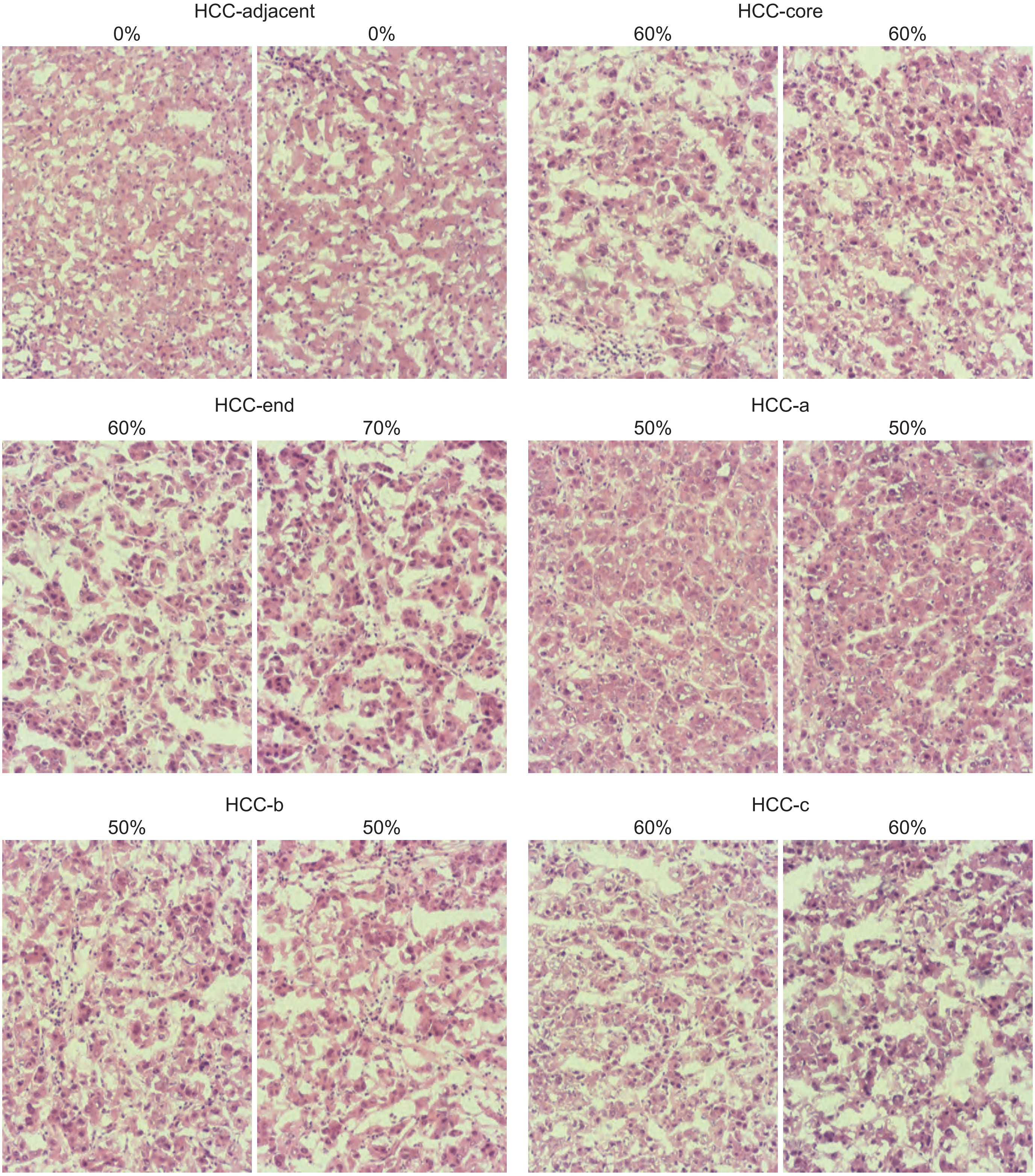


**Supplementary Figure 6.** HE-staining of slices of the ends of five HCC biopsies and of a healthy adjacent biopsy. Tumor percentage (indicated above each picture) was estimated by the pathology department of the UMC Utrecht (the Netherlands) based on these stainings.

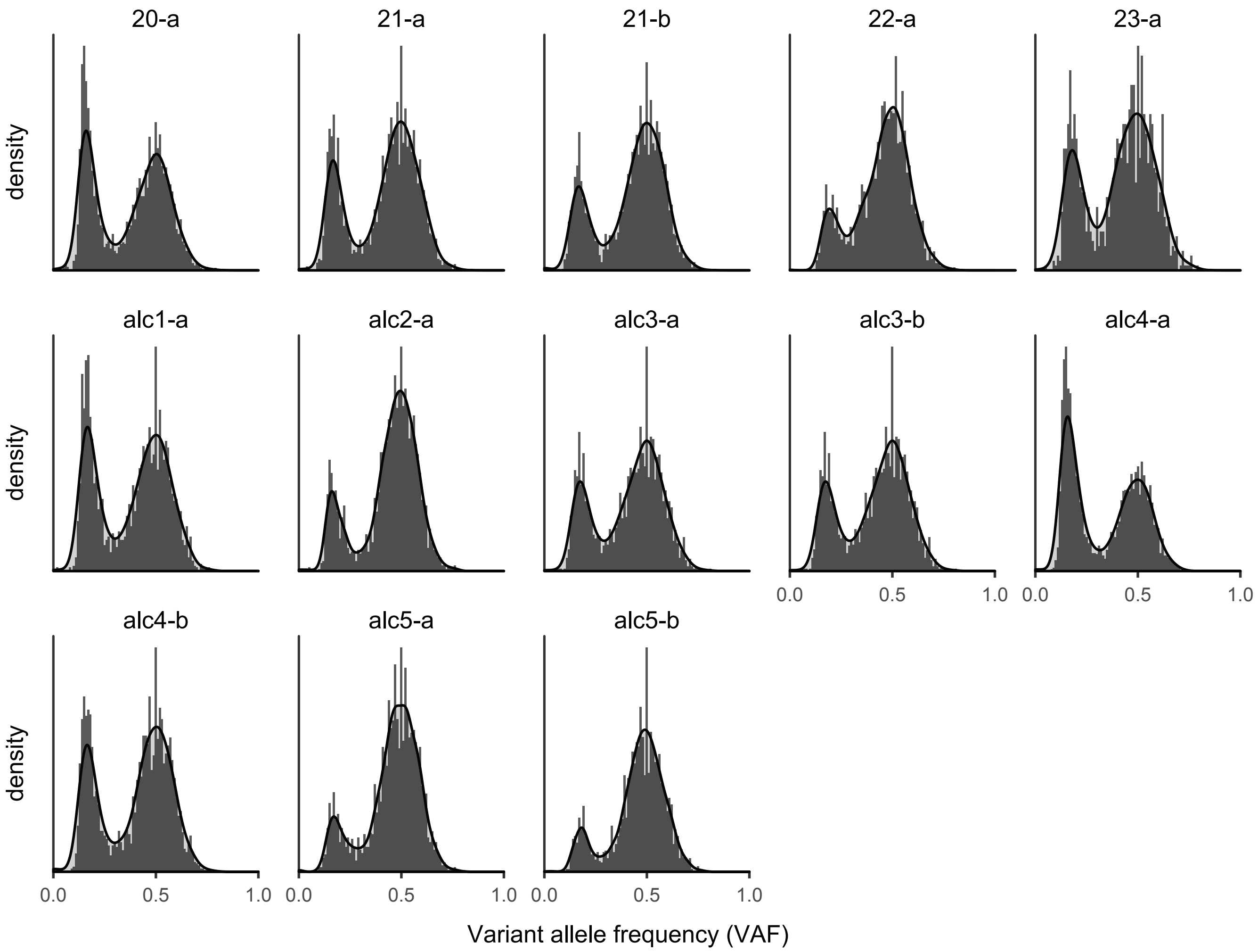


**Supplementary Figure 7.** Variant allele frequency (VAF) distributions of the somatic base substitutions acquired in the genomes of healthy and alcoholic liver ASCs before applying the VAF ≥ 0.3 filter. In this figure, only VAF plots of new samples are displayed. VAF plots of the remaining healthy liver ASCs can be found in Extended Data Figure 2 of Blokzijl *et al.*, [24].

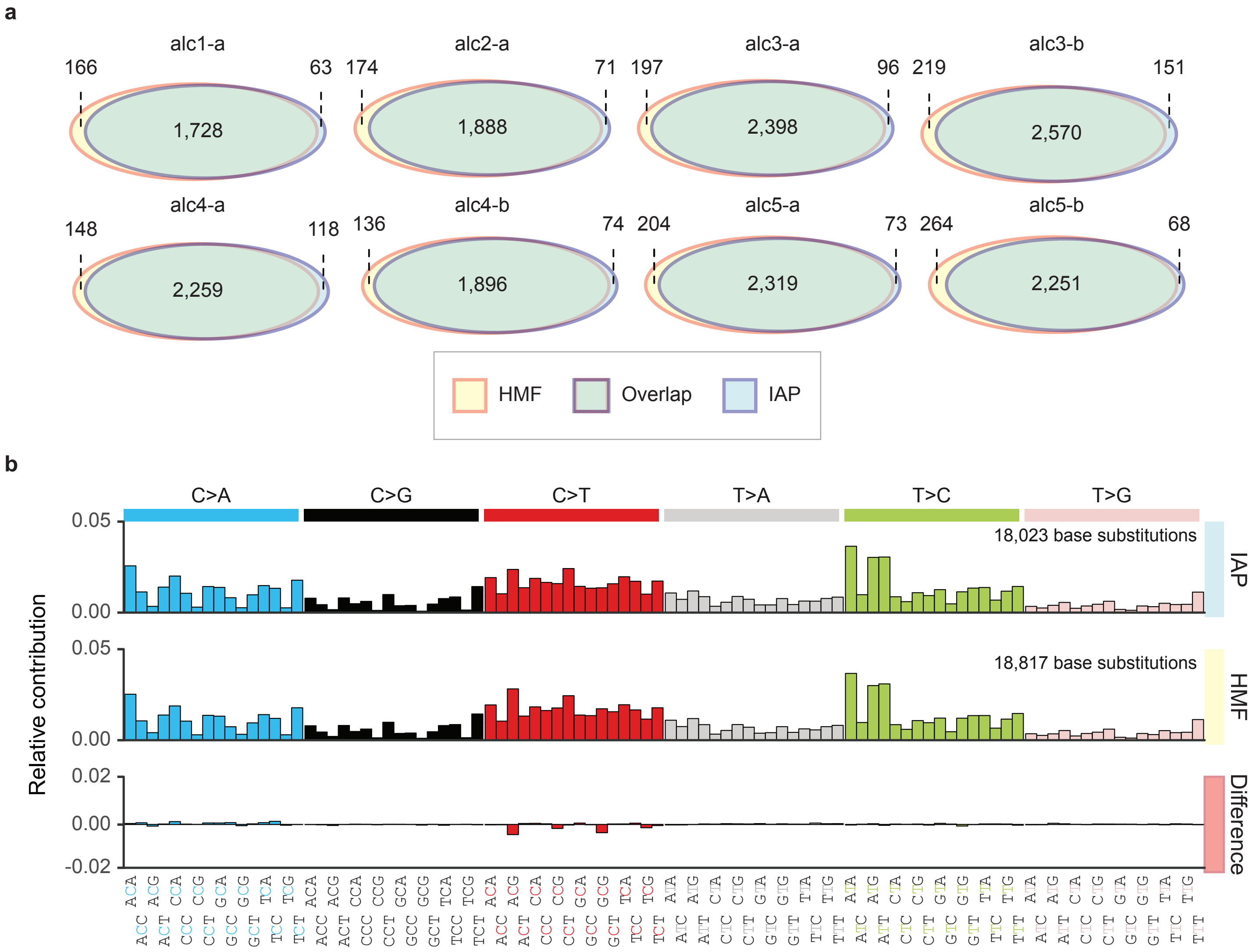


**Supplementary Figure 8.** Comparison between the base substitution catalogs of the alcoholic liver ASCs obtained using two independent variant calling pipelines. The Illumina sequencing pipeline (IAP) (https://github.com/UMCUGenetics/IAP) was used to identify base substitutions in the healthy and alcoholic liver ASCs, whereas the Hartwig Medical Foundation (HMF) pipeline (https://github.com/hartwigmedical) was used to detect base substitutions in the five HCC biopsies. **a** Venn diagrams showing the overlapping variant calls between base substitution catalogs of IAP and HMF per alcoholic liver ASC. A variant was considered to overlap when it was called on the same genomic position. **b** Relative contribution of the 96 context-dependent base substitution types to the mean mutational profiles obtained using IAP and HMF, and to the mean difference between these two profiles.

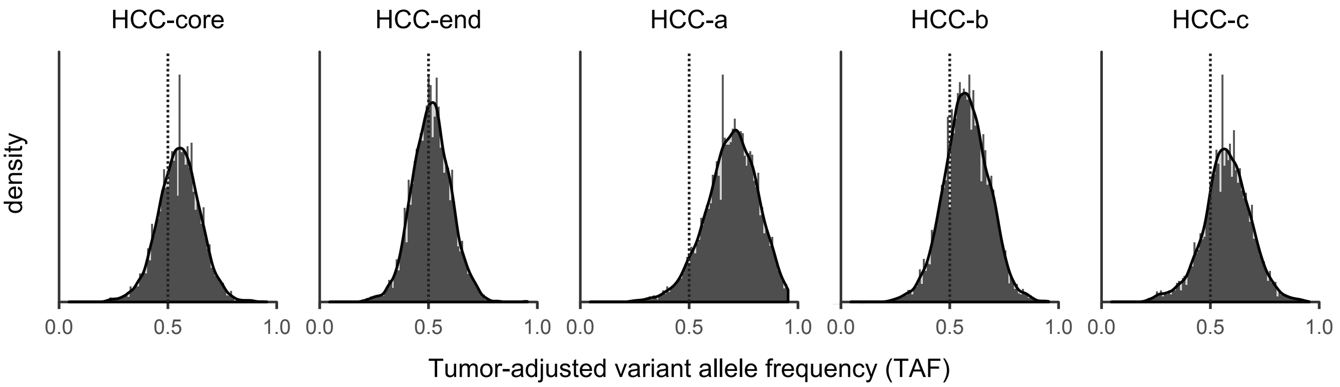


**Supplementary Figure 9.** Distribution of the variant allele frequencies of 6,232 somatic base substitutions that are shared by all five HCC biopsies, per biopsy, adjusted for the estimated tumor percentage per biopsy. Dotted lines indicate a tumor-adjusted variant allele frequency (TAF) of 0.5. Base substitutions on chromosome 1 and 8 were not plotted here, as these chromosomes deviate from a copy number of two in the majority of the biopsies (Supplementary Fig. 4) and the TAF can be affected by this.

| Sample | Donor | Age | Gender | Sample type | Surveyed genome (%) | Base substitutions | Tandem base substitutions | Indels | CNAs* |
|---|---|---|---|---|---|---|---|---|---|
| 23-a | 23 | 24 | Male | Healthy liver | 95.6 | 822 | 2 | 50 | 0 |
| 14-a | 14 | 30 | Male | Healthy liver | 81.4 | 771 | 4 | 18 | 2 |
| 14-b | 14 | 30 | Male | Healthy liver | 85.2 | 888 | 5 | 23 | 0 |
| 15-a | 15 | 41 | Female | Healthy liver | 93.5 | 1,292 | 11 | 48 | 0 |
| 15-b | 15 | 41 | Female | Healthy liver | 95.1 | 1,351 | 8 | 27 | 0 |
| 17-a | 17 | 46 | Female | Healthy liver | 79.4 | 1,495 | 13 | 12 | 0 |
| 17-b | 17 | 46 | Female | Healthy liver | 73.7 | 1,273 | 12 | 12 | 0 |
| 18-c | 18 | 53 | Male | Healthy liver | 97.9 | 1,845 | 10 | 63 | 0 |
| 18-d | 18 | 53 | Male | Healthy liver | 98.5 | 1,504 | 13 | 100 | 1 |
| 18-e | 18 | 53 | Male | Healthy liver | 98.3 | 1,577 | 13 | 89 | 1 |
| 16-a | 16 | 55 | Male | Healthy liver | 97.5 | 1,919 | 13 | 105 | 1 |
| 20-a | 20 | 55 | Female | Healthy liver | 96.2 | 2,066 | 15 | 92 | 0 |
| 22-a | 22 | 61 | Male | Healthy liver | 96.5 | 2,259 | 16 | 130 | 0 |
| 21-a | 21 | 68 | Female | Healthy liver | 96.2 | 2,625 | 19 | 140 | 0 |
| 21-b | 21 | 68 | Female | Healthy liver | 96.5 | 2,383 | 17 | 108 | 0 |
| alc1-a | alc1 | 52 | Male | Alcoholic liver | 96.0 | 1,791 | 16 | 66 | 0 |
| alc2-a | alc2 | 59 | Male | Alcoholic liver | 96.6 | 1,959 | 18 | 104 | 0 |
| alc3-a | alc3 | 60 | Female | Alcoholic liver | 96.0 | 2,494 | 22 | 93 | 0 |
| alc3-b | alc3 | 60 | Female | Alcoholic liver | 96.2 | 2,721 | 24 | 90 | 0 |
| alc4-a | alc4 | 63 | Male | Alcoholic liver | 96.8 | 2,377 | 22 | 114 | 0 |
| alc4-b | alc4 | 63 | Male | Alcoholic liver | 96.7 | 1,970 | 11 | 111 | 0 |
| alc5-a | alc5 | 67 | Male | Alcoholic liver | 96.7 | 2,392 | 20 | 166 | 0 |
| alc5-b | alc5 | 67 | Male | Alcoholic liver | 95.9 | 2,319 | 17 | 170 | 0 |

*CNA = copy number alteration; see extended data table 2 of Blokzijl *et. al* [24] for type and size of CNA

**Supplementary Table 1.** Somatic base substitutions, tandem base substitutions, indels, and copy number alterations acquired in the genomes of healthy and alcoholic liver ASCs.

| Sample | Donor | Age | Gender | Sample type | Base substitutions |
|---|---|---|---|---|---|
| HCC-trunk | HCC1 | 60 | Male | Trunk HCC | 7,203 |
| HCC-core | HCC1 | 60 | Male | HCC biopsy | 9,553 |
| HCC-end | HCC1 | 60 | Male | HCC biopsy | 10,830 |
| HCC-a | HCC1 | 60 | Male | HCC biopsy | 9,347 |
| HCC-b | HCC1 | 60 | Male | HCC biopsy | 10,193 |
| HCC-c | HCC1 | 60 | Male | HCC biopsy | 10,118 |

**Supplementary Table 2.** Somatic base substitutions identified in five biopsies of one alcohol-related HCC.

| Sample | Donor | Age | Gender | Sample type | Chr* | Start | Stop | Size | Type | Genes |
|---|---|---|---|---|---|---|---|---|---|---|
| HCC-trunk | HCC1 | 60 | Male | Trunk HCC | 10 | 60,262,954 | 60,270,868 | 7,915 | Deletion | 0 |
| HCC-trunk | HCC1 | 60 | Male | Trunk HCC | 17 | 2,837,706 | 7,887,745 | 5,050,039 | Deletion | 545 |

* Chr = chromosome

**Supplementary Table 3.** Somatic copy number alterations detected in a recent common ancestor of an alcohol-related HCC.
